# A quantitative model for spatio-temporal dynamics of root gravitropism

**DOI:** 10.1101/2023.04.11.536255

**Authors:** Amir Porat, Mathieu Rivière, Yasmine Meroz

## Abstract

Plant organs adapt their morphology according to environmental signals through growth-driven processes called tropisms. While much effort has been directed in the development of mathematical models describing the tropic dynamics of aerial organs, these cannot provide a good description of roots due to intrinsic physiological differences. Here we present a mathematical model informed by gravitropic experiments on *Arabidopsis thaliana* roots, assuming a sub-apical growth profile and apical sensing. The model quantitatively recovers the full spatio-temporal dynamics observed in experiments. An analytical solution of the model enables us to evaluate the gravitropic and proprioceptive sensitivities of roots, while also allowing us to corroborate the requirement of proprioception in describing root dynamics. Lastly, we find that the dynamics are analogous to a damped harmonic oscillator, providing intuition regarding the source of the observed oscillatory behavior. In all, the model not only provides a quantitative description of root tropic dynamics, but also provides a mathematical framework for the future investigation of roots in complex media.

## Introduction

Plants are sessile organisms, physically anchored to the ground. They survive and prosper in fluctuating environments by adapting their morphology via tropisms the redirection of growth according to external stimuli, such as light and gravity (1–3). As an example, Figs. 1A-B show snapshots during the gravitropic responses of wheat coleoptiles (turning away from gravity), and *Arabidopsis thaliana* roots (growing towards the direction of gravity). The fundamental role of tropisms in the adaptive capabilities of plants has motivated a number of coarse-grained mathematical models describing their macroscopic dynamics (4–10). These models have generally been tailored to describe the dynamics of aerial organs, i.e shoots, since they are experimentally more accessible than roots.

**Fig. 1.**
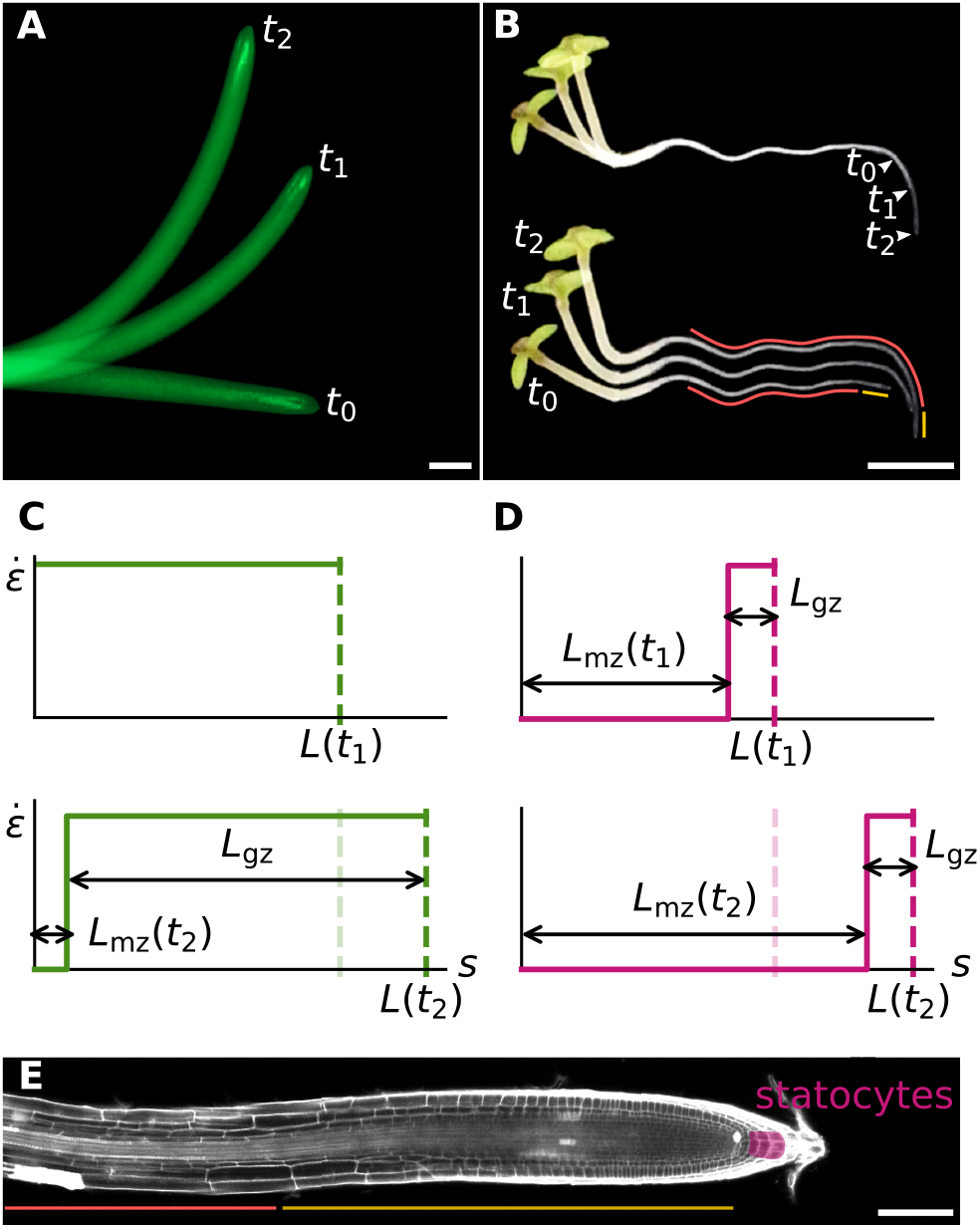
Gravitropism in roots and shoots. **(A)** Negative gravitropic response of a coleoptile of wheat (*Avena sativa*), with snapshots at *t*_0_ = 0, *t*_1_ = 4h, and *t*_2_ = 8h. **(B)** Gravitropic response of an *Arabidopsis thaliana* seedling with snapshots at *t*_0_ = 0, *t*_1_ = 7h, and *t*_2_ = 14h. The hypocotyl (left) displays negative gravitropism while the root (right) shows positive gravitropism. The overlaid picture of the same root at different times (top), demonstrates how the trajectory of the tip matches the final shape of the root. The bottom image, where root images are shown separately, shows how the curvature is produced in the growth zone (yellow line) and progressively fixed in the mature zone (red line). Scale bar: 2 mm. Schematic representation of the spatial distribution of elongation rate along a growing wheat coleoptile at the beginning of gravistimulation (top), and at the end of the experiment (bottom). We mark the length of the whole root *L*(*t*), the mature zone *L*_mz_(*t*), and the growth zone *L*_gz_(*t*) **(D)** Idem for the *Arabidopsis* root. **(E)** Micrograph of an *Arabidopsis thaliana* root. The columella (pink patch) is an order of magnitude smaller than the growth zone (yellow line) and contains inclination-sensitive cells called statocytes that are at the foundation of gravisensing. Scale bar: 100 *μ*m. (Courtesy of Eilon Shani)

However, Figs. 1A-B clearly illustrate that the spatiotemporal dynamics of tropic responses of roots and shoots are very different: While almost the whole coleoptile changes shape in time, in roots most of the shape changes occur close to the tip becoming frozen in time, and the final shape of the root closely matches the trajectory of the tip (11). This is mainly due to physiological differences such as the distribution of the sensory system, and the extent of the growth zone where growth occurs, and curvature is produced as part of tropic responses (6, 12). Figs. 1C-D show a schematic plot of the size and position of the growth zone relative to the organ length, during the gravitropic responses shown in Figs. 1A-B respectively. Most roots grow by several times the length of their growth zone, and therefore most of the produced curvature is frozen in the mature zone (where growth no longer occurs). Conversely shoots grow by only a fraction of length, and changes in shape occur along most of the coleoptile. Furthermore, the distribution of the gravitropic sensory system is different. Gravisensing occurs in specialized sensing cells called statocytes (13–15). In aerial organs gravisensing is local, with statocytes distributed along the whole epidermis in dicots (16, 17), or along the vascular bundle in monocots (16, 18, 19). In roots gravisensing is apical, with statocytes confined to the columella (20) located in the root cap. Fig. 1E illustrates the relative size of the columnella in an *Arabidopsis* root, an order of magnitude smaller than the growth zone.

Here we conduct gravistimulation experiments on *Arabidopsis thaliana* roots, and quantitatively characterize their dynamics. This characterization informs the development of a mathematical model, building on the “angle-curvature-elongation” (ACE) model (6), which accurately describes the observed spatio-temporal dynamics of gravitropic responses of *Arabidopsis* roots.

## Results and Discussion

### Characterization of growth in the gravitropic response of *Arabidopsis* roots

We start with a quantitative characterization of growth as a driver of the tropic response, evaluating the extent of the growth zone relative to the region over which tropic bending occurs. The shape of a root of length *L*(*t*) at time *t* is described by *θ*(*s, t*), the local angle at point *s* along the centerline, where *s* =0 at the base, and *s* = *L*(*t*) at the tip (Fig. 2A). The local curvature is defined as the rate at which the angle changes along the centerline, *k*(*s, t*)= *∂θ*(*s, t*)*/∂s*. The growth zone extends from the tip along a length *L*_gz_. We define *θ*_tip_(*t*)= *θ*(*s* = *L*(*t*), *t*) the angle at the tip, and 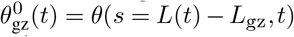 the angle at the base of the growth zone.

**Fig. 2.**
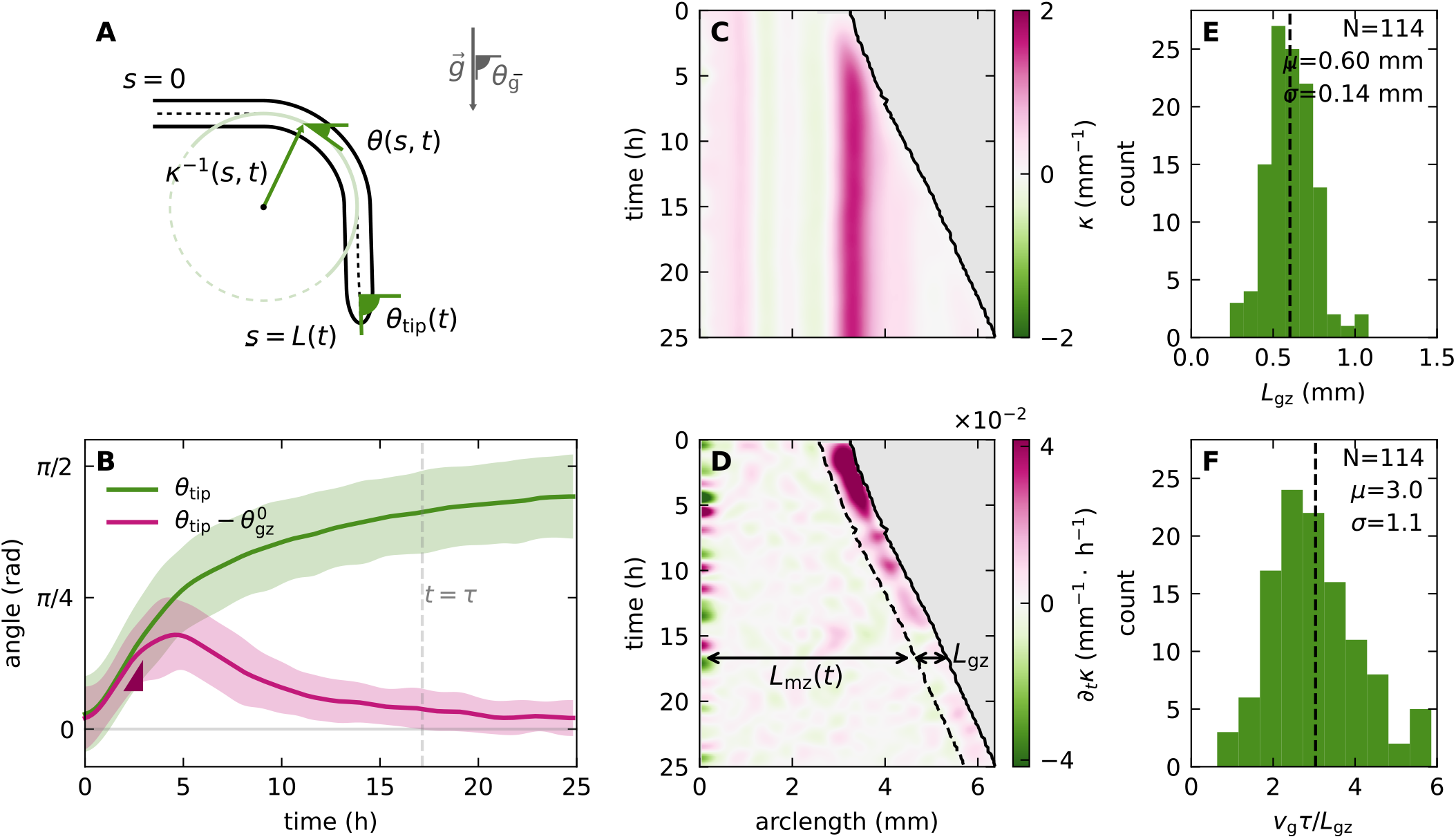
Characterization of tropic responses of Arabidopsis roots. A. **(A)** Geometrical description. *θ*(*s, t*) is the local angle of the root at point *s* along the organ, at time *t*. The local curvature *k*(*s, t*) = *∂θ*(*s, t*)*/∂s* represents how strongly the local angle changes along the organ. *s* = 0 at the base and *s* = *L*(*t*) at the tip, where *L*(*t*) is the organ length, and R is the radius. *θ*_tip_ is the angle at the tip, and *θ*_g_ = *π/*2 is the direction of the gravity signal. **(B)** Average evolution of the tip angle (green) and difference between the angles at the tip and base of the growth zone (pink) during the gravitropic response of *Arabidopsis thaliana* roots tilted horizontally. The vertical dashed line indicates *τ*, the typical time needed for *θ*_tip_ to reach a steady state. **(C)** Graphical representation (kymograph) of the evolution of *k*(*s, t*) along a specific root. **(D)** Kymograph of *∂k*(*s, t*)*/∂t*, highlighting that changes in curvature occur within the growth zone, allowing to assess *L*_gz_ from the observed dynamics. **(E)** Distribution of measured values of *L*_gz_, with an average value of *L*_gz_ = 0.6 mm. **(F)** Distribution of the total growth of roots over their gravitropic response, *L*(*τ*) − *L*_0_ = *v*_g_*τ*, in units of *L*_gz_, where *v*_g_ is the apical growth velocity. On average, *Arabidopsis* roots growth 3 times *L*_gz_ during their gravitropic response, meaning that a mathematical description of the dynamics has to take an explicit account of growth.

These variables provide insight into the dynamics of the gravitropic response (an extended list of the variables used here appears in Table S1). Fig. 2B shows the evolution of the root tip angle *θ*_tip_(*t*) over time, during permanent gravistimulation experiments. Upon being tilted horizontally, the tip angle starts at an angle 0, and increases until it approximately realigns with the direction of gravity. It approaches its steady state value *θ*_f_ *±*σ(*θ*_f_) over a timescale *τ* (see Methods), with the whole growth zone approaching a flat steady state shape, with 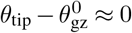.

A full description of the spatio-temporal dynamics is given by the curvature *k*(*s, t*). A typical example is shown in Fig. 2C, showing that *k*(*s, t*) increases within the growth zone near the apex after gravistimulation, and goes back to 0 after a few hours, as manifested in 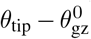. The curvature developed in the growth zone becomes fixed in the mature zone, where growth no longer occurs. This behavior is highlighted further in *k* (*s, t*)*/∂t* (Fig. 2D), where we can identify the growth zone as the region near the tip where changes in curvature occur with *∂k*(*s, t*)*/∂t* ≠ 0. Outside of the growth zone *∂k*(*s, t*)*/∂t* = 0, and the curvature remains constant. This analysis, described in the Methods for multiple roots, enables us to estimate *L*_gz_ for each root and yields an average value of *L*_gz_ = 0.60 *±* 0.14mm (Fig. 2E).

For each root we compare *L*_gz_ to the length grown during the whole gravitropic response (detailed in the Methods section). We find that by the time roots reach their final angle at *t* = *τ*, they grow on average 3 times the growth zone, namely *v*_*g*_*τ* = (3.0 *±*1.1)*L*_gz_ (Fig. 2F), where *v*_*g*_ is the tip growth velocity. This result, in addition to the curvature fixation in the mature zone, confirms the effects of growth cannot be neglected, as can be done in some cases (5).

### The root model

Based on the variables defined before, we build on the ACE model (5, 6) for tropic dynamics with an explicit account of growth. This model has been analyzed and corroborated experimentally for aerial organs (5, 6), and has recently been generalized to 3D (7, 9, 10). The organ length *L*(*t*) evolves according to an axial growth rate profile 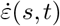, such that the local growth velocity is 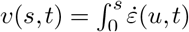. Within the growth zone 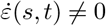, and the general form of the ACE model reads

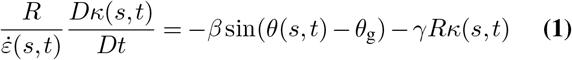

where *θ*_g_ is the direction of the stimulus (here the direction of gravity), *β* is the gravitropic sensitivity or gain, *γ* is the proprioceptive sensitivity, and *D/Dt* = *∂/∂t* + *v*(*s, t*)*∂/∂s* is the material derivative due to growth. The organ is clamped at the base such that *θ*(*s* = 0, *t*)= *θ*_0_, and in the mature zone *Dk*(*s, t*)*/Dt* = 0.

In general we distinguish between two phases of growth during organ development: *Exponential* growth refers to the initial stage where the growth zone spans the length of the whole organ (the case for coleoptiles as illustrated in Fig. 1C, taking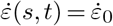 for simplicity), which increases exponentially over time. *Linear* growth refers to the next phase when some tissues stop elongating and *L*(*t*) *> L*_gz_. Only the subapical part of the organ elongates (as for roots, illustrated in Figs. 1D-E), and the total length of the organ increases linearly with time.

As discussed before, in order to describe the dynamics of root growth, one must implement apical sensing and linear growth. Eq. 1 can be modified for apical sensing by replacing *θ*(*s, t*) with *θ*_tip_(*t*), and assuming a constant growth rate 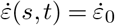 for simplicity, reading in the growth zone

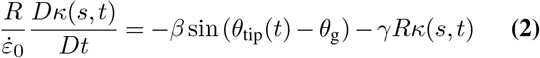

which we will term here the *root model*.

While the steady state shape has been described analytically in the simpler case of exponential growth and local sensing (6), the investigation of linear growth has been limited to numerical studies. Here we focus on a detailed analysis of the root model.

### Solution of the root model

In order to solve Eq. 2, we express the tip angle *θ*_tip_ as a function of curvature, recalling that by definition 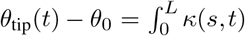. We simplify this relation by distinguishing between the curvature in the growth zone and the mature zone, and splitting the integral accordingly. In the case of apical sensing the signal from the apex is uniform along the growth zone, and assuming an initially straight organ the resulting curvature is also uniform along the growth zone. We therefore rewrite the curvature in the growth zone as a function of time alone *k*(*s, t*)= *k*_gz_(*t*). In the mature zone curvature is fixed, and is therefore a function of arc-length alone, so *k*(*s, t*) = *k*_mz_(*s*). While *L*_gz_ is fixed over time, the mature zone increases. Since the growth velocity is the integral of the growth rate along the growth zone, the organ length increases linearly according to 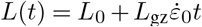, where *L*_0_ is the initial length of the organ such that *L*_0_ *> L*_gz_. The length of the mature zone is then

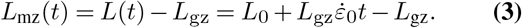

We now simplify the expression for *θ*_tip_(*t*) by splitting the integral over the GZ and MZ, and substituting *k*_gz_(*t*) and *k*_mz_(*s*) accordingly:

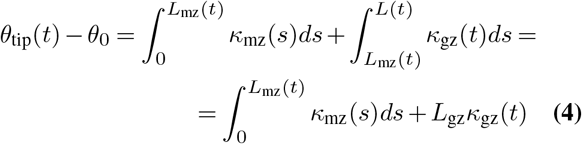

To continue, we note that the curvature at a point *s* in the mature zone *k*_mz_(*s*) was fixed when it exited the growth zone at some previous time *t*^*′*^ *< t*, which allows to relate it to a past value of the curvature of the growth zone *k*_mz_(*s*) = *k*_gz_(*t*^*′*^). Since the basal end of the growth zone is equivalent to the length of the mature zone, the time *t*^*′*^ at which arc length *s* left the growth zone can be found by *s* = *L*_mz_(*t*^*′*^). Assuming for simplicity that *L*_0_ = *L*_gz_, Eq. 3 yields 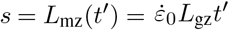, which allows us to change variables in the last integral in Eq. 4 from *s* to *t*^*′*^:

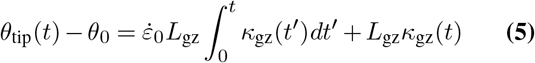

We substitute Eq. 5 into Eq. 2. The material derivative now becomes a simple time derivative since *k*_gz_(*t*) is as a function of time alone. Together, the PDE of the original ACE model is now transformed to an ordinary integro-differential equation for *k*_gz_(*t*):

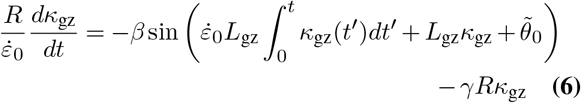

where 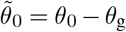 is the initial angle relative to the gravitational stimulus.

We rewrite Eq. 6 in a non-dimensional form: (i) normalizing curvature according to *L*_gz_, defining k = L_gz_*k*_gz_ equivalent to the angle traced by the growth zone 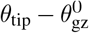, and (ii) normalizing time according to the growth rate, defining 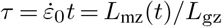 equivalent to the organ growth in units of *L*_gz_. Lastly, we define *η* = *βL*_gz_*/R* (experimental values listed in Table 1), where *L*_gz_*/R* is the geometric slenderness ratio of the growth zone, and *η* acts as an effective gravitropic sensitivity. Substituting these non-dimensional quantities in Eq. 6, and denoting *d*k*/*d*τ* ≡ k*′*, yields

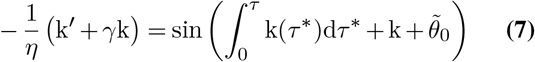

Applying arcsin and taking another derivative in *τ* results in

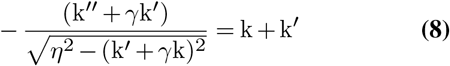

Eq. 8 is a second order non-linear ODE for k(*τ*) with the initial conditions k(0) = 0 and 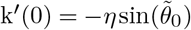. We note that a straight organ, with k(*τ*)= 0, is a steady state solution. This equation can be easily solved numerically, and we find the evolution of *θ*_tip_(*t*) by substituting the solution of Eq. 8 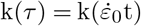in the normalized form of Eq. 5:

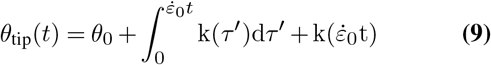

**Table 1.**
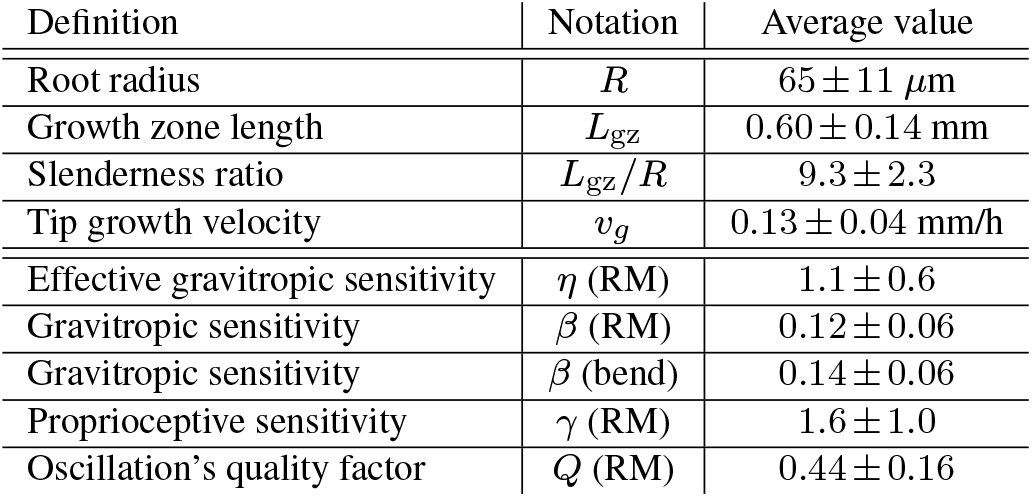
Measured slenderness ratios and root model (RM) fitted variables with *R*^2^ *>* 0.9 (88 out of 114 roots)

| Definition | Notation | Average value |
| --- | --- | --- |
| Root radius | $R$ | $65 \pm 11 \mu\text{m}$ |
| Growth zone length | $L_{gz}$ | $0.60 \pm 0.14 \text{ mm}$ |
| Slenderness ratio | $L_{gz}/R$ | $9.3 \pm 2.3$ |
| Tip growth velocity | $v_g$ | $0.13 \pm 0.04 \text{ mm/h}$ |
| Effective gravitropic sensitivity | $\eta$ (RM) | $1.1 \pm 0.6$ |
| Gravitropic sensitivity | $\beta$ (RM) | $0.12 \pm 0.06$ |
| Gravitropic sensitivity | $\beta$ (bend) | $0.14 \pm 0.06$ |
| Proprioceptive sensitivity | $\gamma$ (RM) | $1.6 \pm 1.0$ |
| Oscillation's quality factor | $Q$ (RM) | $0.44 \pm 0.16$ |

### Qualitative analysis of root model solutions and comparison to other models

Fig. 3A shows the evolution of *θ*_tip_(*t*) solved for the gravitropic response of a root tilted horizontally, with 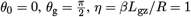 (based on typical values for *Arabidopsis* found in the literature (21), see Table 1), and for a range of proprioception values *γ*. In all cases the tip eventually aligns with the direction of gravity, and the averaged experimental trajectory follows a similar trend. We later quantitatively discuss the fit of experimental trajectories with root model solutions. For comparison, Figs. 3B-C show the corresponding simulated trajectories for the AC model, which does not account for growth, and the ACE model with exponential growth, both with apical sensing. In the SM we develop the analytical solutions plotted here for both models. In the AC model the tip angle never aligns with the direction of gravity apart from the case with no prioprioception *γ* = 0, and by definition it does not account for growth, describing only cases with *L*(*t*)*/L*_gz_ ≪1.

**Fig. 3.**
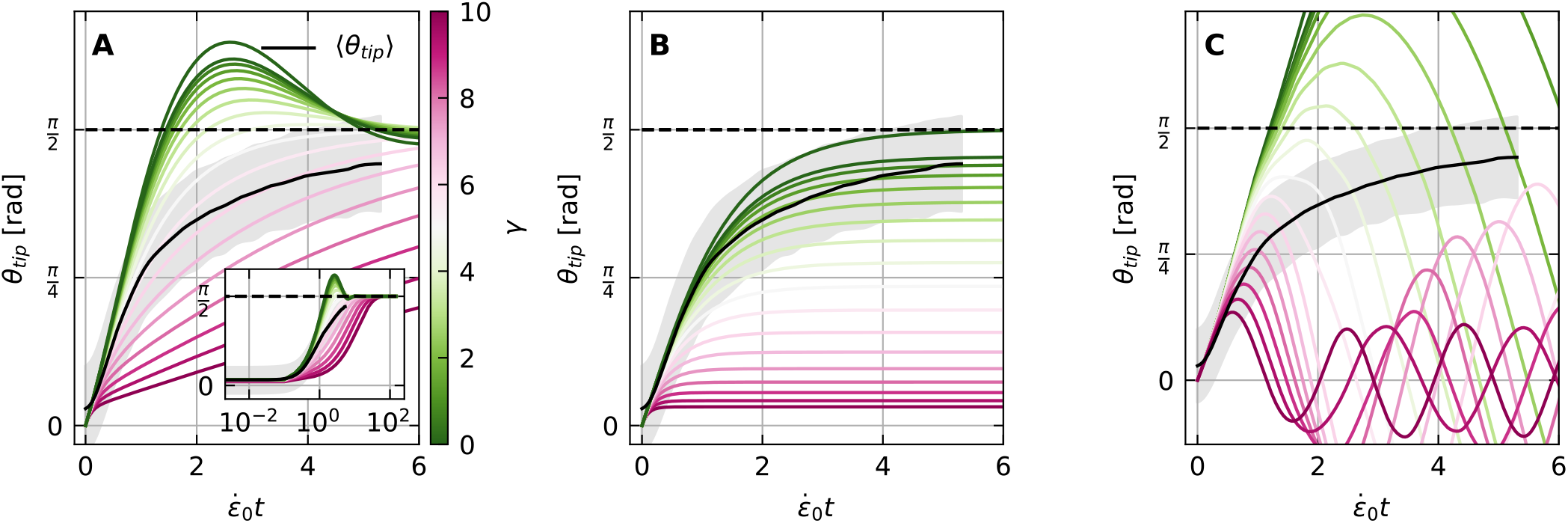
Simulations of gravitropic responses of *Arabidopsis* roots for different models. **(A)** Root model. Simulated evolution of *θ*_tip_ solving the root model, obtained by numerically solving Eq. 8, and substituting the solution into Eq. 9. Here *θ*_g_ = *π/*2, *θ*_0_ = 0 and *η* = *βL*_*gz*_*/R* = 1, for proprioceptive values *γ* ranging 0 to 10. Over long times the tip angle always reaches the direction of gravity *θ*_g_ = *π/*2 marked by a dashed line, as also found in experiments in Fig. 2, here plotted by a black line. **(B)** AC model. Evolution of *θ*_tip_ for the AC model (5), where growth is implicitly assumed to be the driver of the tropic response, but not taken into account explicitly. Simulations use the same parameters as in (A). The tip aligns with the stimulus only in the case of *γ* = 0. **(C)** ACE model with exponential growth. Evolution of *θ*_tip_ for the ACE model (6), where the growth zone is assumed to include the whole organ. Simulations use the same parameters as in (A) and (B). The tip overshoots the direction of the stimulus in all cases.

We also recall that in the case of apical sensing the AC model leads to a constant curvature throughout the root organ, which, as we discuss later, does not reflect the spatial growth pattern of roots. Despite these inconsistencies the AC model manages to fit the tip angle trajectories well, and in the SM we compare its fitting results to those of the root model. Lastly, the trajectories of the ACE model with exponential growth exhibit significant oscillations, overshooting the direction of gravity and never reaching a steady state. The deviation from experimental trajectories is such that attempts to fit to the exponential ACE model do not converge.

### Fitting tip angle trajectories and parameter estimation

We fit *θ*_tip_(*t*) trajectories of the gravitropic experiments to numerical solutions of the root model, Eqs. 8 and 9, detailed in the Methods. The fitting parameters are the proprioceptive sensitivity *γ*, the effective gravitropic sensitivity *η* = *βL*_gz_*/R*, and the initial angle *θ*_0_. The fit therefore provides an estimation of these parameters, and 77% of experiments fit with *R*^2^ *>* 0.9 (Fig. S1). The rest of the parameters, *R, v*_*g*_ and *L*_gz_, are extracted directly through image analysis, while 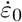 is estimated *via v*_*g*_*/L*_gz_ (see Methods). Parameters are summarized in Table 1.

We compare the values of *β* (Fig. 4A) estimated for single trajectories using the root model, to an alternative estimation method based on the maximal bending of the growth zone (14):

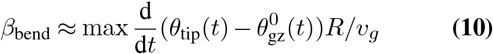

where *v*_*g*_ is the tip growth velocity (slope shown in Fig. 2B). The two methods agree on the average gravitropic sensitivity, with average values ⟨*β*_RM_ ⟩= 0.12 0.06 and ⟨*β*_bend_ ⟩= 0.14 *±*0.06 respectively.

**Fig. 4.**
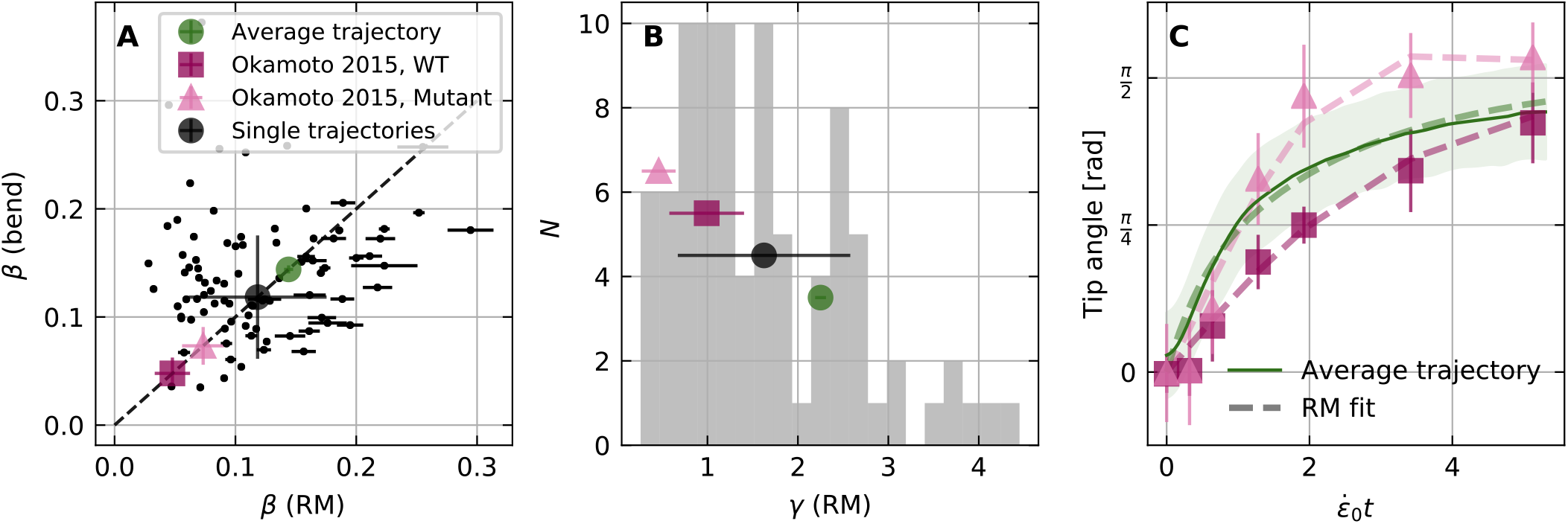
Estimation of gravitropic and proprioceptive sensitivities *β* and *γ* by fitting tip angle trajectories to root model. **(A)** The gravitropic sensitivity *β* estimated using two methods: (i) fitting the *θ*_tip_ trajectories measured for single *Arabidopsis* roots with solutions of the root model from Eq. 9 (with *R*^2^ *>* 0.9, see Fig. S1), which gives *β* = 0.12 *±* 0.06 and (ii) extracted by the maximal initial bending (Eq. 10) which gives *β* = 0.14 *±* 0.06. Single trajectories are plotted (black dots) along with their ensemble statistics (large black circle). **(B)** Histogram (shaded area) of the proprioceptive coefficient values *γ* estimated by fitting *θ*_tip_ trajectories to solutions of the root model (with *R*^2^ *>* 0.9, see Fig. S1). The average value is *γ* = 1.6 *±* 1.0 (black circle). **(C)** Fit of averaged trajectories using the root model solutions for three different data sets: (i) solid green line represents average trajectory of our data, dashed line represents the fitted root model solution with *β* = 0.14 *±* 0.00 and *γ* = 2.3 *±* 0.1, (ii) purple squares represent average trajectories for WT from (22), and the dashed line represents the root model fit with *β* = 0.05 *±* 0.01 and *γ* = 1.0 *±* 0.4, and (iii) pink triangles represents hyper-bending mutants from (22), and the dashed line represents the rot model fit with *β* = 0.07 *±* 0.02 and *γ* = 0.5 *±* 0.2. Colored symbols representing the fitted *β* and *γ* appear also in (A) and (B).

This further corroborates the root model, and we now proceed to evaluate the proprioceptive coefficient *γ* – the first systematic estimation for roots. Fig. 4B presents the distribution of the values of *γ* estimated by fitting the root model for single *θ*_tip_ trajectories, with an average value ⟨*γ ⟩* = 1.6 *±*1.0. We attempted to fit the trajectories to a modified root model with *γ* = 0. Only 5% of trajectories fit well with *R*^2^ *>* 0.9 (Fig. S1), confirming that proprioception is required in the description of root growth dynamics. Furthermore, we analyzed data from Okamoto et al. (22) for ACTIN8, an *Arabidopsis* mutant defective in myosin XI exhibiting hyper bending in its gravitropic response, and predicted to correspond to a lower proprioceptive gain. We fit the averaged *θ*_tip_ trajectories for 35 WT roots and 35 mutant roots digitized from *Okamoto et al*. (22), as well as the average trajectory from Fig. 2, using the root model (Fig. 4C). Estimated values of *β* and *γ* are summarized in Table 2 and Fig. 5 (colored symbols), where as predicted the proprioceptive value for the overshooting mutant is lower than for WT. As values for *v*_*g*_, *R* and *L*_gz_ were not provided, and we used our values instead (Table 1), the direct comparison between values extracted from *Okamoto et al*. and our own data is limited.

**Table 2.** Estimations of *β* and *γ* based on the root model using data from Okamoto et al. 2015 (22). The hyper-bending mutants were fitted with a lower proprioceptive coefficient than the WT.

| | $\beta$ | $\gamma$ |
| --- | --- | --- |
| WT (Okamoto et al. 2015) | $0.05 \pm 0.01$ | $1.0 \pm 0.4$ |
| Mutant (Okamoto et al. 2015) | $0.07 \pm 0.02$ | $0.5 \pm 0.2$ |

**Fig. 5.**
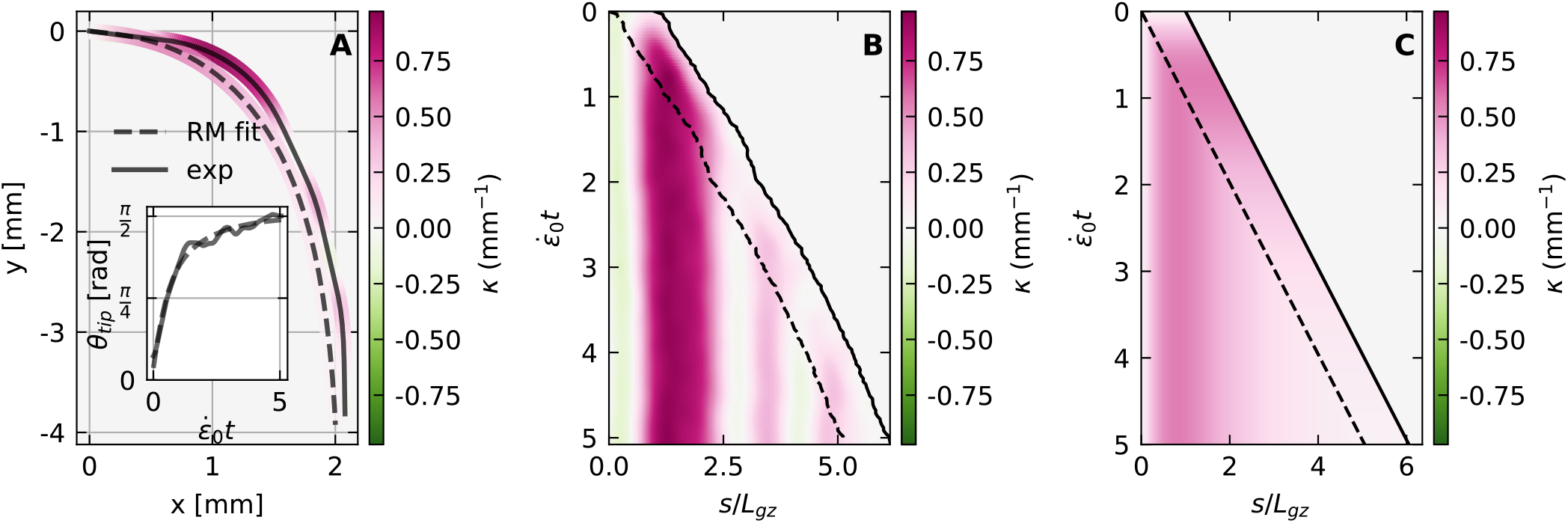
Root model recovers full spatio-temporal dynamics of organ shape. **(A)** Example of a specific *Arabidopsis* root shape (solid line) overlayed with the simulated shape using the parameters extracted from a fit to the *θ*_tip_ trajectory using the solutions of the root model. The color represents curvature, as in the kymographs in B and C. The inset shows the corresponding *θ*_tip_ trajectory (solid black) along with its fit (dashed line) based on Eq. 9 with a coefficient of determination *R*^2^ = 0.98. The fitted sensitivities are *γ* = 1.6 *±* 0.0 and *β* = 0.17 *±* 0.00. **(B)** Graphical representation (kymograph) of the evolution of *k*(*s, t*) along a specific root, which tip angle and shape are presented in **A**. The constant growth zone *L*_gz_ and increasing mature zone *L*_mz_ are delineated with a dashed line. **(C)** The simulated kymograph based on the root model fit.

### Root model recovers full spatio-temporal dynamics of organ

The tip angle dynamics provides limited insight into the shape of the whole organ throughout the response. For example, while the AC model with apical sensing provides a good description of *θ*_tip_ trajectories (Fig. S1), it wrongly predicts a uniform curvature throughout the organ (Fig. 2B). We analyze and compare the full spatio-temporal dynamics of roots and the root model. Fig. 5A shows an example of the final shape of a specific root, where colors represent the local curvature *k*(*s*). The dashed line represents the simulated root, based on the parameters estimated from the fit to *θ*_tip_ of the same root (shown in the inset), in agreement with the observed form. Fig. 5B presents a kymograph of the measured *k*(*s, t*) of the root over time, and Fig. 5C shows the simulated *k*(*s, t*) of the same root. In order to compare experiment with simulation, we normalize time with 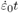 and space with *s/L*_gz_. The growth-zone *L*_gz_ is marked on both figures with a dashed line. We find that the simulation captures the main features of the spatio-temporal dynamics, namely the initial production of curvature in the growth zone at short times, which is then fixed at the base of the organ (11). We note that despite the similarity with the simulated root, the curvature in the growth zone of actual roots is not uniform, as illustrated and quantified in Figs. S2,S3.

### Small angle approximation provides an analogy with a damped harmonic oscillator

In order to gain further insight, we analyze the root model in the limit of a small angle approximation |*θ*_tip_(*t*) − *θ*_g_ | ≪1. This is the variable of the sine function in Eq. 7, enabling us to linearize the equation derivative in *τ* yields such that 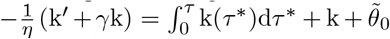. Taking a derivative in *τ* yields

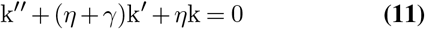

This can be identified as the equation for a damped harmonic oscillator, and the analogy provides an intuitive understanding of the dynamics. We identify two normalized time-scales: (i) the natural angular frequency 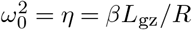, associated with the natural oscillations with no damping, and (ii) the attenuation rate r = (*η* + *γ*), associated with damping due to friction. Damped oscillations, corresponding to roots overshooting the direction of gravity, occur when 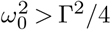, i.e. 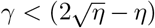. The quality factor is defined as 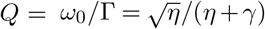, and describes how quickly oscillations die out. We note that the oscillations timescale related to the natural angular frequency, 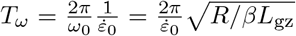, is proportional to the related timescale *T*_*f*_ estimated in (6) for the non-apical ACE model; the time required for a an element aligned with the direction of gravity to leave the growth zone and become fixed.

We can identify the source of the oscillations as the term *η*k in Eq. 11, which can be traced back to the integral over the curvature exiting the growth zone in Eq. 4. We note that neglecting the corresponding integral in Eq. 6 recovers the AC model with no growth. Indeed, the tip angle in the apical AC model does not present oscillations and decays exponentially to its steady state angle with an attenuation rate equivalent to *η* + *γ* (Fig. 3). This destabilizing source of oscillations corresponds to *passive orientation drift* (6), where the tip angle of a growing organ with a fixed curvature increases, even with no differential growth. This drift leads to an overshoot of the direction of gravity, as clearly seen in Fig.3A for low values of *γ*.

## Materials and Methods

### Plant material and growth conditions

The wild-type strain of *Arabidopsis thaliana* was used in this study. Sterile seeds were plated on a growth medium consisting of half-strength Murashige and Skoog medium adjusted to pH 5.7, 1% sucrose and 1% phytagel. Seeds were stratified for 2 days at 4^°^*C* and then kept vertical in a growth chamber at 24^°^*C* under a 12 h photoperiod.

### Experimental protocol

After germination, seedlings were transferred to another Petri dish under a biological hood. Seedlings were arranged by groups of five and the roots were placed between the lid of the Petri dish and a block of fresh phytagel (same content as the growth medium). This new Petri dish was then mounted on a rotating plate. During the experiment, the plate was maintained vertical for 10 h, then tilted horizontally for the following 24 h. White light was provided from the top throughout the experiment, except when taking pictures (light from all directions). Pictures were taken every 10 min using a Nikon D750. The inclination angle, lighting and camera were all controlled by an Arduino micro-controller running a specifically developed program.

### Image analysis

The noise and contrast of the acquired pictures were edited using the free software RawTherapee (http://rawtherapee.com/) to ease further analysis steps. We used a version of Interekt (23) to extract the shape of each root at each time step of the experiment, yielding the root arc length *s*, its local angle with respect to the horizontal *θ*(*s, t*), the total length *L*(*t*), and the radius *R*(*s, t*) (see Fig. 1F). The radius was averaged over both *s* and *t*. Assuming linear growth, the tip growth velocity *v*_*g*_ = *v*(*s* = *L*(*t*), *t*) was extracted as the slope of a linear fit of *L*(*t*). We defined three further angles: (i) the tip angle *θ*_tip_(*t*) *θ*(*s* = *L*(*t*), *t*) and (ii) the angle at the base of the growth zone, at the interface of the mature zone, 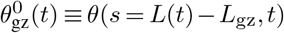. Both were obtained by locally averaging *θ*(*s, t*) over a characteristic length of 0.1 mm around the points of interest. Lastly, (iii) the tip angle in the final steady state *θ*_f_, defined as the average of *θ*_tip_(*t*) over the last 5h of the experiment, and σ(*θ*_f_) the corresponding standard deviation. The curvature *k*(*s, t*) was calculated according to *k* = *∂θ/∂s*. We define the time to reach gravitropic equilibrium *τ* as the time for *θ*_tip_(*t*) to approach its steady state value *θ*_f_ *±*σ(*θ*_f_) Assuming that growth is the only driver of curvature changes as in the ACE model, we estimated the growth zone *L*_gz_ indirectly by looking at extent over which curvature changes over time *∂k*(*s, t*)*/∂t* = 0, as clearly seen in Fig. 2D. We estimate *L*_gz_ from 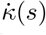 the spatial profile of *∂k/∂t* in the frame attached to the tip and averaged in time for *t< τ*. *L*_gz_ is then defined as the minimal distance from the tip such that 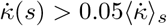, where ⟨· ⟩_*s*_ denotes averaging over *s*. The growth rate in the growth zone is approximated according to 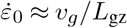.

### Curve fitting

The fitting of experimental trajectories of *θ*_tip_(*t*) to Eq. 9 and the second order ordinary differential equation (ODE) in Eq. 8 was done using the `curve_fit` function from the open source SciPy Python library. This function called the `odeint` function from the same library, which solved the ODE multiple times using the free fittingparameters {*γ, η, θ*} until the fitting was complete. Lastly, since both 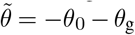 and 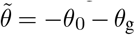 yield the same initial condition for 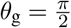, we bounded the fitting of *θ*_0_ using *θ*_0_ *>* 0.

## Conclusions

Here we present the root model, a robust mathematical description of the spatio-temporal dynamics of plant root gravitropic responses. The model is based on physiological differences between roots and shoots informed by experiments on *Arabidopsis thaliana*, namely apical sensing and a finite growth zone resulting in linear growth.

We show that the root model captures the dynamics of the tip angle, as well as the full spatio-temporal dynamics of the root’s gravitropic response over time. Furthermore, we find an analytical solution to the model, allowing to estimate the gravitropic and proprioceptive sensitivities, and we find that proprioception is necessary in order to describe root responses. This first dynamical quantification of proprioception in plant roots corroborates mutants associated with a lower proprioceptive gain (22).

In the limit of a small angle approximation, the root model is analogous to a damped harmonic oscillator, providing intuition for the observed dynamics. The observed damped oscillatory dynamics can be expressed directly using the quality factor of the root dynamics which has the low value of *Q* = 0.44 *±*0.16.

The model discussed here focuses on gravitropism, however, as discussed elsewhere (10), it can be easily generalized to take into account other types of tropisms, such as hydrotropism and thigmotropism, thus providing a theoretical framework for the quantitative study of root growth in complex media.

## ACKNOWLEDGEMENTS

The authors are grateful to Odelia Pisanty, Hamutal Bar and Eilon Shani for providing the seeds as well as for their help and guidance in the cultivation of the plants. AP would like to thank Ilana Taieb Hagerty and Roni Kempinski for the fruitful discussions.

## AUTHOR CONTRIBUTIONS

Formal analysis, Software, Methodology, AP and MR; Data curation, MR; Conceptualization and Investigation, AP MR and YM; Visualization, Writing – Original draft, Review and editing, AP MR and YM; Funding acquisition, Supervision, Resources, Project admnistration YM.

## DATA AVAILABILITY

Experimental trajectories available at: mathieuriviere/Porat2023_RootModel.git https://github.com/

## Bibliography

1. Simon Gilroy and Patrick Masson. Plant Tropisms. Wiley-Blackwell, 2007.

2. Lucius Wilhelminus Franciscus Muthert, Luigi Gennaro Izzo, Martijn van Zanten, and Giovanna Aronne. Root tropisms: Investigations on earth and in space to unravel plant growth direction. Frontiers in Plant Science, 10:1807, 2020. ISSN 1664-462X. doi: 10.3389/fpls.2019.01807.

3. Yasmine Meroz. Plant tropisms as a window on plant computational processes. New Phytologist, 229(4):1911–1916, 2021.

4. Thomas Guillon, Yves Dumont, and Thierry Fourcaud. A new mathematical framework for modelling the biomechanics of growing trees with rod theory. Mathematical and Computer Modelling, 55(9-10):2061–2077, 2012.

5. Renaud Bastien, Tomas Bohr, Bruno Moulia, and Stéphane Douady. Unifying model of shoot gravitropism reveals proprioception as a central feature of posture control in plants. Proceedings of the National Academy of Sciences of the United States of America, 110: 755–760, January 2013.

6. Renaud Bastien, S. phane Douady, and Bruno Moulia. A unifying modeling of plant shoot gravitropism with an explicit account of the effects of growth. Frontiers in Plant Science, 5, April 2014.

7. Daniele Agostinelli, Alessandro Lucantonio, Giovanni Noselli, and Antonio DeSimone. Nutations in growing plant shoots: The role of elastic deformations due to gravity loading. Journal of the Mechanics and Physics of Solids, 136:103702, 2020. ISSN 0022-5096. doi: https://doi.org/10.1016/j.jmps.2019.103702. The Davide Bigoni 60th Anniversary Issue.

8. Alberto Bressan, Michele Palladino, and Wen Shen. Growth models for tree stems and vines. Journal of Differential Equations, 263(4):2280–2316, 2017.

9. Derek E. Moulton, Hadrien Oliveri, and Alain Goriely. Multiscale integration of environmental stimuli in plant tropism produces complex behaviors. Proceedings of the National Academy of Sciences, 117(51):32226–32237, 2020. ISSN 0027-8424. doi: 10.1073/pnas.2016025117.

10. Amir Porat, Fabio Tedone, Michele Palladino, Pierangelo Marcati, and Yasmine Meroz. A general 3d model for growth dynamics of sensory-growth systems: From plants to robotics. Frontiers in Robotics and AI, 7:89, 2020. ISSN 2296-9144. doi: 10.3389/frobt.2020.00089.

11. Andrés Chavarría-Krauser, Kerstin A Nagel, Klaus Palme, Ulrich Schurr, Achim Walter, and Hanno Scharr. Spatio-temporal quantification of differential growth processes in root growth zones based on a novel combination of image sequence processing and refined concepts describing curvature production. New Phytologist, 177(3):811–821, 2008.

12. Renaud Bastien, Stéphane Douady, and Bruno Moulia. A Unified Model of Shoot Tropism in Plants: Photo-, Gravi- and Propio-ception. PLoS Comput Biol, 11(2):e1004037, February 2015.

13. Yasuko Hashiguchi, Masao Tasaka, and Miyo T Morita. Mechanism of higher plant gravity sensing. American Journal of Botany, 100(1):91–100, 2013.

14. Hugo Chauvet, Olivier Pouliquen, Yoël Forterre, Valérie Legué, and Bruno Moulia. Inclination not force is sensed by plants during shoot gravitropism. Scientific reports, 6(1):1–8, 2016.

15. Olivier Pouliquen, Yoël Forterre Antoine Bérut, Hugo Chauvet, François Bizet, Valérie Legué, and Bruno Moulia. A new scenario for gravity detection in plants: the position sensor hypothesis. Physical Biology, 14(3):035005, 2017.

16. Lilian E Hawker. A quantitative study of the geotropism of seedlings with special reference to the nature and development of their statolith apparatus. Annals of Botany, 46(181):121–157, 1932.

17. Kenichiro Fujihira, Tetsuya Kurata, Masaaki K Watahiki, Ichirou Karahara, and Kotaro T Yamamoto. An agravitropic mutant of arabidopsis, endodermal-amyloplast less 1, that lacks amyloplasts in hypocotyl endodermal cell layer. Plant and Cell Physiology, 41(11):1193–1199, 2000.

18. FD Sack, D. Priestley, and AC Leopold. Surface charge on isolated maize-coleoptile amyloplasts. Planta, 157(6):511–517, 1983.

19. U Kutschera. Gravitropism of axial organs in multicellular plants. Advances in space research, 27(5):851–860, 2001.

20. Elison B Blancaflor, Jeremiah M Fasano, and Simon Gilroy. Mapping the functional roles of cap cells in the response of arabidopsis primary roots to gravity. Plant physiology, 116(1):213–222, 1998.

21. J Roué, H Chauvet, N Brunel-Michac, F Bizet, B Moulia, E Badel, and V Legué. Root cap size and shape influence responses to the physical strength of the growth medium in Arabidopsis thaliana primary roots. Journal of Experimental Botany, 71(1):126–137,11 2019. ISSN 0022-0957. doi: 10.1093/jxb/erz418.

22. Keishi Okamoto, Haruko Ueda, Tomoo Shimada, Kentaro Tamura, Takehide Kato, Masao Tasaka, Miyo Terao Morita, and Ikuko Hara-Nishimura. Regulation of organ straightening and plant posture by an actin–myosin xi cytoskeleton. Nature plants, 1(4):1–7, 2015.

23. Félix P Hartmann, Hugo Chauvet-Thiry, Jérôme Franchel, Stéphane Ploquin, Bruno Moulia, Nathalie Leblanc-Fournier, and Mélanie Decourteix. Methods for a quantitative comparison of gravitropism and posture control over a wide range of herbaceous and woody species. In Plant Gravitropism, pages 117–131. Springer, 2022.

