## Supplementary_material for "A quantitative model for spatio-temporal dynamics of root gravitropism"

### Supplementary Note 1: Tip angles solutions for the AC and the exponential ACE models

Here we find an expression for $\theta_{\text{tip}}(t)$ for the AC model and ACE model with exponential growth, in the case of apical sensing, solving the dynamical equations. These expressions are plotted in Fig. 3B-C.

***AC model with apical sensing.*** The AC model with apical sensing follows (5)

$$\frac{d\kappa(t)}{dt} = \frac{\dot{\varepsilon}_0}{R}\left(-\beta\sin(\theta_{\text{tip}}(t)-\theta_{\text{g}}) - \gamma R\kappa(t)\right), \tag{S1}$$

for $s$ in the growth zone. We note that the model is written in this form in order to properly compare it to the root model. In the AC model growth is not considered explicitly, and the length of the organ is constant $L(t) = L_0$. In the case of apical sensing, an initially straight organ will develop constant curvature throughout the growth zone, and since $\theta_{\text{tip}} - \theta_0 = \int_0^{L_0} \kappa ds$, we have

$$\theta_{\text{tip}} - \theta_0 = \kappa L_0 \tag{S2}$$

Substituting Eq. S2 in Eq. S1, and integrating along $s$ following Bastien et al. (12) yields

$$\frac{1}{\dot{\varepsilon}_0}\frac{d\theta_{\text{tip}}}{dt} = -\frac{\beta L_0}{R}\sin(\theta_{\text{tip}}-\theta_{\text{g}}) - \gamma(\theta_{\text{tip}}-\theta_0). \tag{S3}$$

We take $\theta_{\text{g}} = \pi/2$, and the dynamics of $\theta_{\text{tip}}(t)$ can be found by

$$\dot{\varepsilon}_0 t = \int_{\theta_0}^{\theta_{\text{tip}}(t)} \frac{d\theta_{\text{tip}}}{\frac{\beta L_0}{R}\cos(\theta_{\text{tip}}) - \gamma(\theta_{\text{tip}}-\theta_0)} \tag{S4}$$

We note that the time diverges if $(\theta_{\text{tip}}-\theta_0) = \beta L_0\cos(\theta_{\text{tip}})/(R\gamma)$, giving an expression for the steady state tip angle. When $\gamma = 0$, we obtain

$$\theta_{\text{tip}}(t) = \arcsin\left(\tanh\left(\frac{\beta L_0}{R}\dot{\varepsilon}t + c\right)\right) \tag{S5}$$

where $c = \text{artanh}(\sin(\theta_0))$. In the small angle approximation, Eq. S3 can be written as

$$\frac{1}{\dot{\varepsilon}_0}\frac{d\theta_{\text{tip}}}{dt} = -\left(\frac{\beta L_0}{R} + \gamma\right)\theta_{\text{tip}} + \frac{\beta L_0}{R}\theta_{\text{g}} + \gamma\theta_0 \tag{S6}$$

Such that the damping rate of the dynamics is $\beta L_0/R + \gamma$, similar to the non-dimensional rate $\eta + \gamma$ found in the main text for the root model. Lastly, in order to fit the tip angle trajectories to the dynamics of the AC model, we use a non-dimensional version of Eq. S3 by denoting $\tau = \dot{\varepsilon}_0 t$, $\frac{d\theta_{\text{tip}}}{d\tau} \equiv \theta'_{\text{tip}}$ and $\eta = \beta L_0/R$

$$\theta'_{\text{tip}} = -\eta\sin(\theta_{\text{tip}}-\theta_{\text{g}}) - \gamma(\theta_{\text{tip}}-\theta_0) \tag{S7}$$

Rearranging Eq. S8, taking arcsin on both sides and then taking another derivative of $\tau$ yields

$$-\frac{(\theta''_{\text{tip}} + \gamma\theta'_{\text{tip}})}{\sqrt{\eta^2 - (\theta'_{\text{tip}} + \gamma(\theta_{\text{tip}}-\theta_0))^2}} = \theta'_{\text{tip}} \tag{S8}$$

We then fitted the tip angle trajectories to Eq. S8 as described in the methods for the root model. Note the similarity between Eq. S8 and Eq. 8. We plot $\theta_{\text{tip}}$ for different values of $\gamma$ in In Fig. 3B, and compare between the fitted values of $\gamma$ and $\beta$ of the AC model and the root model in Fig. S1.

***ACE model with exponential growth and apical sensing.*** Assuming a constant growth rate along the organ, we have $\frac{dL(t)}{dt} = \dot{\varepsilon}_0 L(t)$, leading to exponential growth

$$L(t) = L_0 e^{\dot{\varepsilon}_0 t} \tag{S9}$$

As before, curvature is constant for apical sensing, and the tip angle follows

$$\theta_{\text{tip}}(t) - \theta_0 = L_0 e^{\dot{\varepsilon}_0 t}\kappa(t). \tag{S10}$$

Substituting this expression in the ACE model in Eq. 2 gives

$$\frac{d\kappa}{dt} = -\frac{\dot{\varepsilon}_0}{R}\left(\beta\sin(\theta_0 - \theta_{\text{g}} + L_0 e^{\dot{\varepsilon}_0 t}\kappa) + \gamma R\kappa\right). \tag{S11}$$

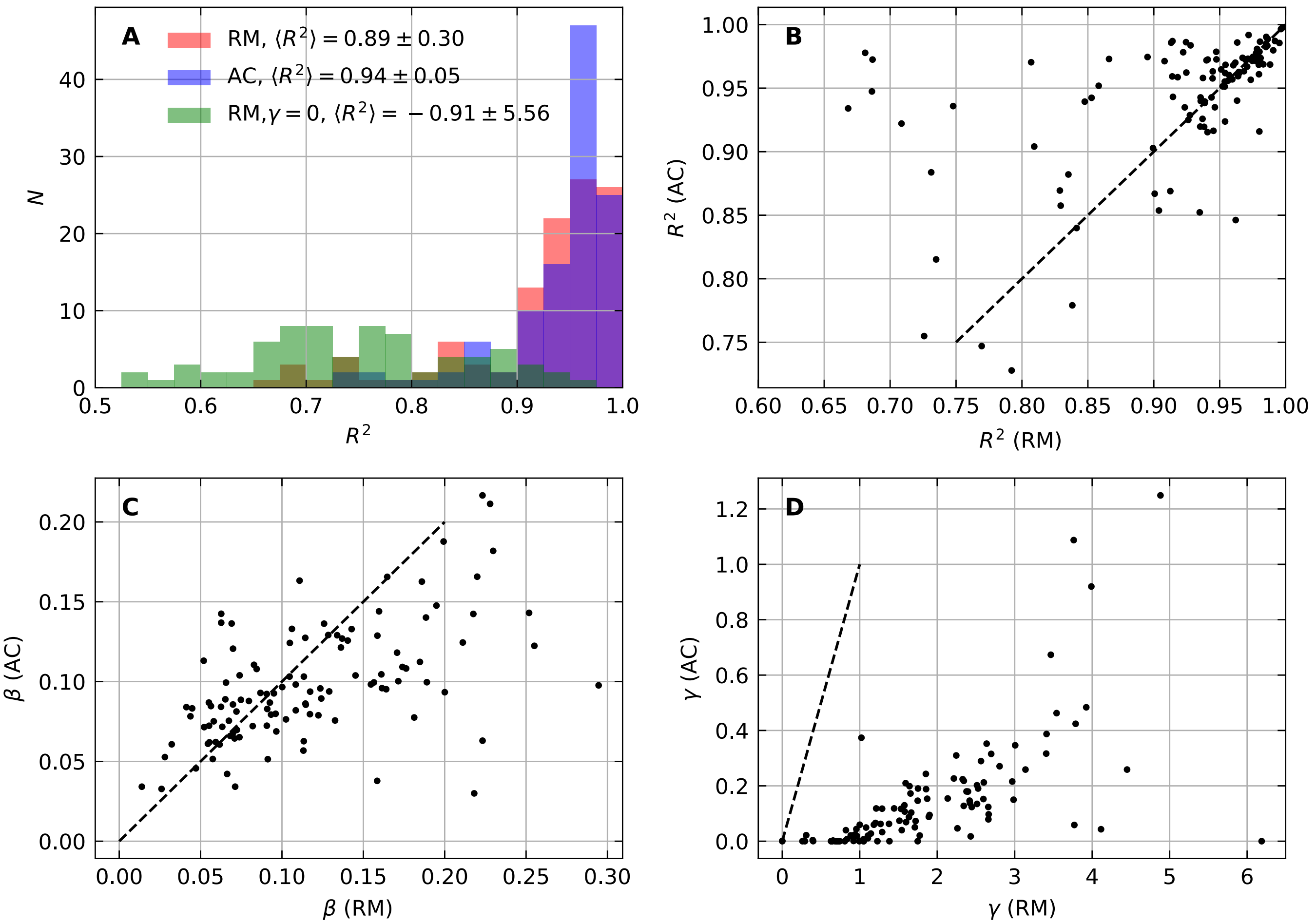


**Fig. S1. Comparison of parameters extracted from the root model and the AC model. (A)** Histograms of the coefficient of determination $R^2$ regarding the fitting of tip angle trajectories using the root model (RM), the root model without proprioception ($\gamma = 0$) and the AC model. **(B)** Comparison between $R^2$ estimated by the root model and the apical AC model. The dashed line is the identity function. **(C)** Comparison between $\beta$ estimated by the root model and the apical AC model. The dashed line is the identity function. **(D)** Comparison between $\gamma$ estimated by the root model and the apical AC model. The dashed line is the identity function.

Rewriting this expression in a non-dimensional form, substituting $\mathrm{k} = \mathrm{L}_0\kappa$, $\tau = \dot{\varepsilon}_0 t$ and $\eta = \beta L_0/R$ as in the main text gives

$$\frac{d\mathrm{k}}{d\tau} = -\eta \sin\left(\theta_0 - \theta_\mathrm{g} + e^{\tau}\mathrm{k}\right) - \gamma \mathrm{k}. \tag{S12}$$

This is an ODE with an initial condition $\mathrm{k}(0) = 0$. We note that k never reaches a steady state, and the angle oscillates at all times as shown in (6). In Fig. 3C we integrate it by assuming $\theta_0 - \theta_\mathrm{g} = -\pi/2$. Fitting of the experimental data to Eq. S12 did not converge using standard fitting methods as described in the methods section.

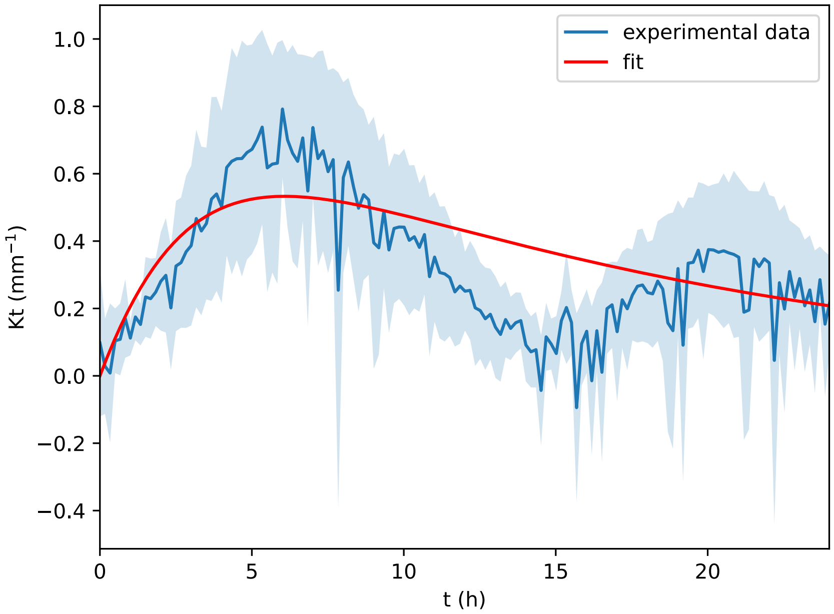


**Fig. S2.** Comparison of the experimental $\kappa_{\mathrm{gz}}$ to the fitted one. The dark blue line is the average measured curvature in the growth zone, and the light blue describes its standard deviation.

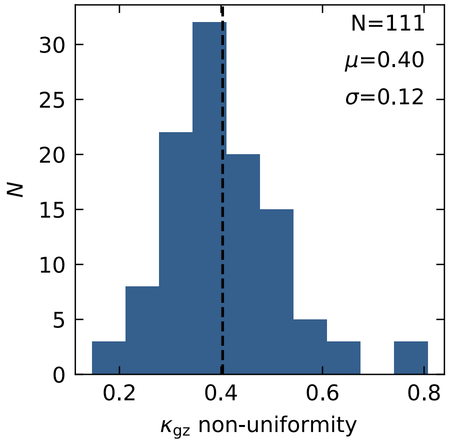


**Fig. S3. Measure of the non-uniformity of curvature within the growth zone.** The non-uniformity was defined as the noise to signal ratio $\sigma(\kappa_{\mathrm{gz}}(t))/\kappa_{\mathrm{gz}}(t)$ averaged over times $t$ such that $\kappa_{\mathrm{gz}}(t) \geq \kappa^{\mathrm{f}}_{\mathrm{gz}} + 2\sigma(\kappa^{\mathrm{f}}_{\mathrm{gz}})$, where $\kappa^{\mathrm{f}}_{\mathrm{gz}} = \langle \kappa_{\mathrm{gz}}(t) \rangle_{t \geq \tau}$

| Notation | Definition | Mathematical definition |
|---|---|---|
| $t$ | time | |
| $s$ | arc length | |
| $v(s,t)$ | axial growth velocity | |
| $v_g$ | tip growth velocity | $v(s=L(t),t)$ |
| $L(t)$ | total length | |
| $L_{\mathrm{gz}}$ | growth zone length | |
| $L_0$ | initial length | |
| $L_{\mathrm{mz}}(t)$ | mature zone length | $L(t)-L_{\mathrm{gz}}$ |
| $R$ | root radius | |
| $\theta(s,t)$ | local angle | |
| $\theta_{\mathrm{tip}}(t)$ | angle at the apex | $\theta(s=L(t),t)$ |
| $\theta_{\mathrm{gz}}^{0}(t)$ | angle at the base of the growth zone | $\theta(s=L(t)-L_{\mathrm{gz}},t)$ |
| $\theta_{\mathrm{f}}$ | steady-state angle | $\langle\theta_{\mathrm{tip}}(t)\rangle_{t\geq 20h}$ |
| $\theta_{\mathrm{g}}$ | angle of the stimulation | |
| $\theta_0$ | initial angle of the root | |
| $\kappa(s,t)$ | curvature | $\partial\theta(s,t)/\partial s$ |
| $\kappa_{\mathrm{gz}}(t)$ | curvature in the growth zone | $\langle\kappa(s,t)\rangle_{s\in GZ}$ |
| $\kappa_{\mathrm{mz}}(s)$ | curvature in the mature zone | |
| $\kappa_{\mathrm{gz}}^{\mathrm{f}}$ | steady-state value of $\kappa_{\mathrm{gz}}$ | $\langle\kappa_{\mathrm{gz}}(t)\rangle_{t\geq\tau}$ |
| $\tau$ | time to reach $\theta_{\mathrm{f}}\pm\sigma(\theta_{\mathrm{f}})$ | |
| $\dot{\varepsilon}_0$ | elongation rate in growth zone | |
| $\beta$ | gravitropic sensitivity | |
| $\gamma$ | proprioceptive sensitivity | |
| k | non-dimensional curvature | $L_{\mathrm{gz}}\kappa_{\mathrm{gz}}(t)$ |
| $\tau$ | non-dimensional time | $\dot{\varepsilon}_0 t$ |
| $\eta$ | effective gravitropic sensitivity | $\beta L_{\mathrm{gz}}/R$ |

**Table S1.** Notations
